# Aging clocks, entropy, and the limits of age-reversal

**DOI:** 10.1101/2022.02.06.479300

**Authors:** Andrei E. Tarkhov, Kirill A. Denisov, Peter O. Fedichev

## Abstract

We analyze aging signatures of DNA methylation and longitudinal electronic medical records from the UK Biobank datasets and observe that aging is driven by a large number of independent and infrequent transitions between metastable states in a vast configuration space. The compound effect of configuration changes can be captured by a single stochastic variable, thermodynamic biological age (tBA), tracking entropy produced, and hence information lost during aging. We show that tBA increases with age, causes the linear and irreversible drift of physiological state variables, reduces resilience, and drives the exponential acceleration of chronic disease incidence and death risks. The entropic character of aging drift sets severe constraints on the possibilities of age reversal. However, we highlight the universal features of configuration transitions, suggest practical ways of suppressing the rate of aging in humans, and speculate on the possibility of achieving negligible senescence.

## I. INTRODUCTION

Aging is a complex process manifesting itself across different organismal levels (see hallmarks of aging [1]) and leading to the exponential acceleration of the incidence of chronic diseases [2] and mortality [3]. It is both practically and intellectually appealing to reduce the effects of the multitude of phenotypic changes to a few, or, even better, a single actionable indicator, most commonly referred to as “biological age”. Biological age (BA) models can be trained either to predict chronological age or mortality risks of an individual from different sources of biomedical data, ranging from DNA methylation (DNAm) [4–13] to physical activity records from wearable devices [14, 15]. Excessive BA (or biological age acceleration) is associated with all-cause mortality, the prevalence, future incidence, and severity of chronic [9, 16, 17] and transient diseases, including COVID-19 [15, 18–20]. Hence, BA predictors have increasingly gained traction in clinical trials [21–23].

The dynamic properties of BA and the exact relation between BA variation and aging are not entirely understood. For example, epigenetic aging drift and hence methylation age may increase without an appreciable increase in all-cause mortality in negligible senescent species [24, 25]. Moreover, even in the most healthy individuals, BA levels can transiently change throughout the day following circadian rhythms [26] or in response to stress factors and lifestyle choices such as smoking [17, 27]. The characteristic time required for an organism state to relax toward the homeostatic equilibrium and the range of BA fluctuations progressively increase as a function of age [17]. The number of individuals exhibiting slow recovery increases exponentially and doubles approximately every 8 years, which is close to the mortality doubling time for humans [15]. Further applications of BA models in aging research and medicine would require a better understanding of the dynamics and causal relation between, on the one hand, underlying biological and physiological variations of the organism state captured by various BA indicators and, on the other, mortality, the prevalence and severity of diseases and the effects of medical interventions.

To address these fundamental questions, we reviewed the universal features of aging signatures in biomedical data. We performed the principal component analysis (PCA) in a large cross-sectional white-blood-cells DNA methylation (DNAm) dataset [28] and in the longitudinal electronic medical records (EMRs) from the UK Biobank [29]. In both cases, we observed a large number of infrequent transitions between the respective states representing the methylation of individual CpG sites or the incidence of specific diseases in the course of aging. At the same time, most of the variance in the data could be explained by the stochastic evolution of a single factor linearly increasing with age and demonstrating the strongest correlation with Horvath’s DNAm age and the number of chronic diseases in the DNAm and EMR datasets, respectively.

To explain the dynamics behind the universally observed aging signatures, we put forward a semiquantitative model of aging in a complex regulatory network. We assumed that living systems are collections of a vast number of interacting functional units (FUs), each set to a metastable state at the end of development. Aging then results from the stochastic relaxation of the organism state towards equilibrium via a sequence of configuration transitions representing the microscopic state changes in all FUs.

Since the number of dynamically accessible irregular configurations is vast, the total number of configuration transitions is also large. Hence, their compound effects on individual biological processes can be quantified by a stochastic variable with a linearly increasing mean and variance. The quantity progressively increases over time in a sufficiently large regulatory network and hence may provide a natural aging clock – the thermodynamic biological age (tBA). We argue that tBA is the fundamental aging variable best associated with the dominant principal component (PC) score in biomedical data, the Horvath methylation clock, and is proportional to the configuration entropy and hence quantifies the information lost in the course of the aging process.

## II. RESULTS

### A. Aging signatures in cross-sectional DNAm data

We start by analyzing a DNAm dataset GSE87571 [28] produced from aging leukocytes measured by the Illumina Infinium 450K Human Methylation Beadchip. Each of the reported DNAm levels 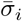 is the average of a binary single-cell signal over a bulk tissue sample comprising a large number of cells. In other words, 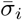 is the probability of finding a CpG site in a methylated state.

To avoid complications due to the cross-over between the development and aging process, we removed samples collected from individuals younger than 25 y.o. Furthermore, to counter the “curse of dimensionality” [30] due to the shallow nature of the dataset (approximately 450k CpGs, each measured in less than 800 patients), we filtered out CpGs without a significant correlation with age (after the Bonferroni correction for multiple testing, *p* = 0.005*/*450k). Ouf of approximately 100k CpGs significantly correlated with age (almost 25% of all reported), most were either initially hypermethylated in the 20 − 25-year-olds (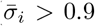, 26% of all CpGs) or hypomethylated (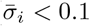, 28% of all CpGs).

The DNAm signal has a non-normal distribution because it is limited to the interval [0, 1]. Instead, we used normally-distributed log-odds ratios 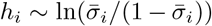) characterizing the probability of a CpG’s methylation. In physics, such a parametrization is used to define the probability of observing a polarized state of a magnet in the presence of a magnetic field according to the Boltzmann probability distribution (see Section S.V). Accordingly, *h*_*i*_ quantifies the free energy differences between the corresponding methylation states. In view of this analogy, henceforth, we refer to *h*_*i*_ as “regulatory fields”.

The PCA of regulatory fields reveals a few principal components (PCs) associated with age. The dominant PC (DNAm-PC1) evolves approximately linearly as a function of age (Fig. 1a; Pearson’s *r* = 0.68, *p* = 3 ·10^−98^). The variance of DNAm-PC1 also increases linearly with age (Fig. 1b), which is a hallmark of a stochastic process (see below).

**FIG. 1.**
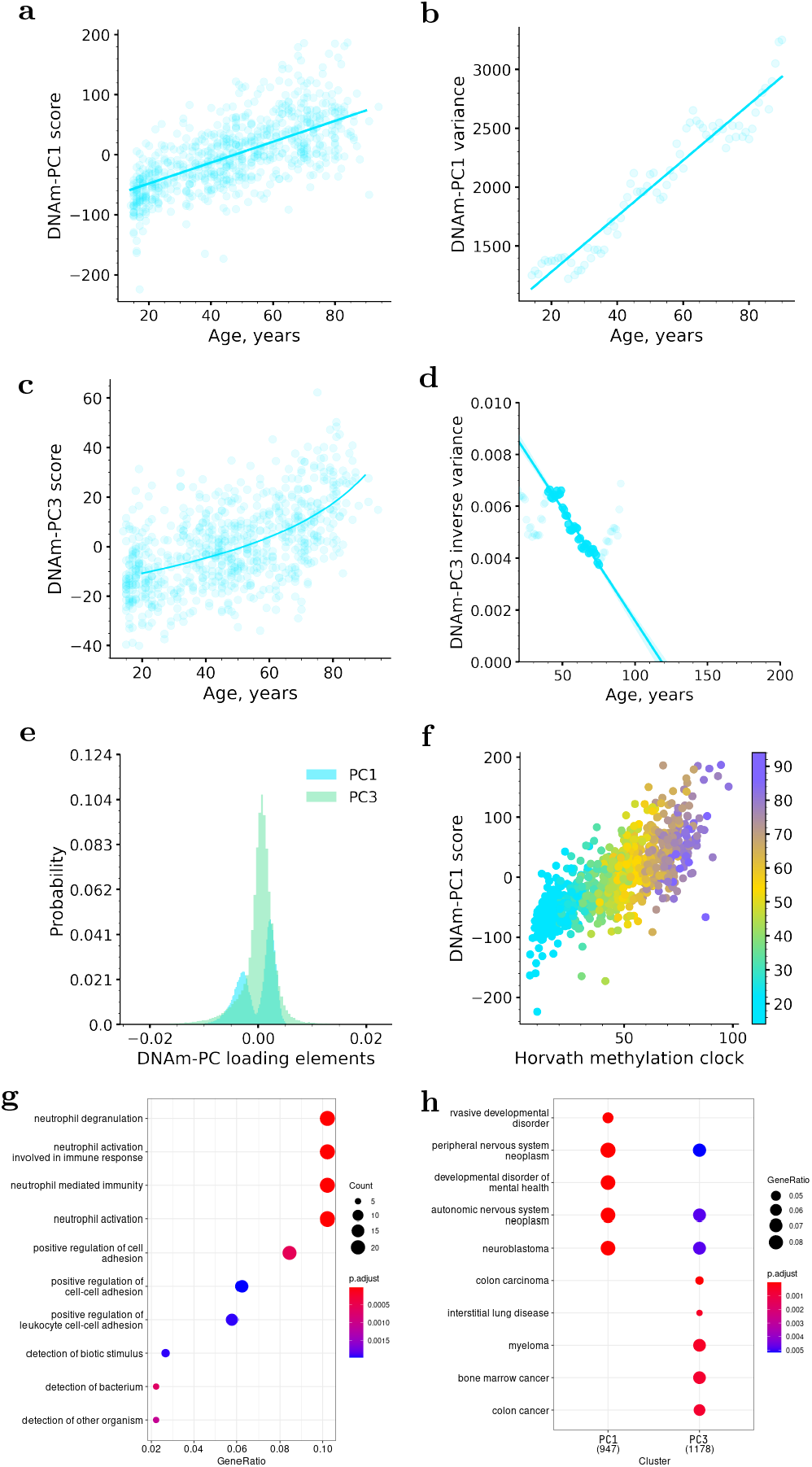
PCA of age-dependent methylation profiles from white-blood-cells samples from GSE87571. (a) DNAm-PC1 and (b) variance of DNAm-PC1 increase, on average, linearly as functions of age. (c) DNAm-PC3 increases faster than linearly with age; (d) inverse variance of DNAm-PC3, the extrapolation for the range of 40 to 75 years vanishes at 120 y.o. (e) Distribution of PC vector components. (f) DNAm-PC1 scores correlate with Horvath’s DNAm age. The color bar represents patients’ chronological age. (g) Gene set enrichment analysis: CpG sites comprising DNAm-PC3 are associated with the regulation of innate immune response. (h) Methylation profiles driven by DNAm-PC1 and DNAm-PC3 are associated with developmental and mental diseases, and internal organs’ diseases.

Aside from DNAm-PC1, the best correlation with chronological age was produced by the third PC, DNAm-PC3, (Pearson’s *r* = 0.56, *p* = 3 ·10^−62^). DNAm-PC3 in subsequent age-adjusted bins increased faster than at a linear pace as a function of age (Fig. 1c). The variance of DNAm-PC3 also increased faster than linearly so that the inverse variance decreased approximately linearly in the patients older than 40 years old (Fig. 1d). By extrapolation, the inverse variance of DNAm-PC3 would approach zero (and hence the variance would diverge) at some age within the age range of 120–150 years.

The loading vectors corresponding to DNAm-PC1 and -PC3 describe two distinct methylation profile changes with age. The distribution of the PC1 loading vector’s components is non-Gaussian and bi-modal. Hence, the dominant aging signature in DNAm data involves two large groups of CpG sites (Fig. 1e) changing their methylation (“polarization”) with age in opposite directions. The first PC score is then proportional to the total number of polarization transitions.

On the contrary, the distribution of the loading vector’s components from DNAm-PC3 has a single peak and clear leading contributions from non-Gaussian tails (see Fig. 1e). The gene set enrichment analysis of methylation regions associated with the PC3 variation reveals pathways involved in innate immunity and cancer (Figs. 1g and h).

The dominant PC scores computed from the GSE87571 samples demonstrate the best correlation with Horvarth’s DNAm age [4] (Fig. 1f). The corresponding Pearson’s correlation coefficients were *r* = 0.75 (*p* = 2 · 10^−131^) and *r* = 0.52 (*p* = 10^−52^) for DNAm-PC1 and DNAm-PC3, respectively (see also Figs. S.1 and S.2 for a summary of other PC scores’ correlation with age and Horvath’s DNAm age).

To confirm the stochastic character of the dominant aging signature in humans, one would need to analyze a large longitudinal dataset. We did not have access to a high-quality set of longitudinal DNAm measurements. Instead, we turned to an extensive electronic medical records collection (EMRs) from the UK Biobank. Irrespective of the age at the first assessment, the EMRs provided information on the prevalence of chronic dis-eases from birth until the end of the follow-up (slightly more than ten years after enrollment, on average). We represent each patient by a vector of binary variables indicating the presence or absence of a disease (see Section S.III).

Most of UK Biobank’s subjects are healthy early in life. Hence, the states representing the presence of diseases are initially polarized (*σ*_*i*_ = 0). Most states stay polarized for life, with only a small fraction of patients exhibiting depolarization transitions leading to the incidence of specific diseases. Indeed, chronic diseases are relatively infrequent: the most prevalent diseases are metabolic disorders (with the prevalence of 15%), joint disorders (14%), and arthrosis (12%).

The PCA of binary-valued vectors representing a health state for the EMRs of UKB subjects at the time of the first assessment look similar to the PCA results from the white-blood-cells DNAm study above. This time, we observed only two PCs significantly associated with age (see the blue and the green lines and the respective ranges corresponding to the mean levels and one standard deviation in Fig. 2a).

**FIG. 2.**
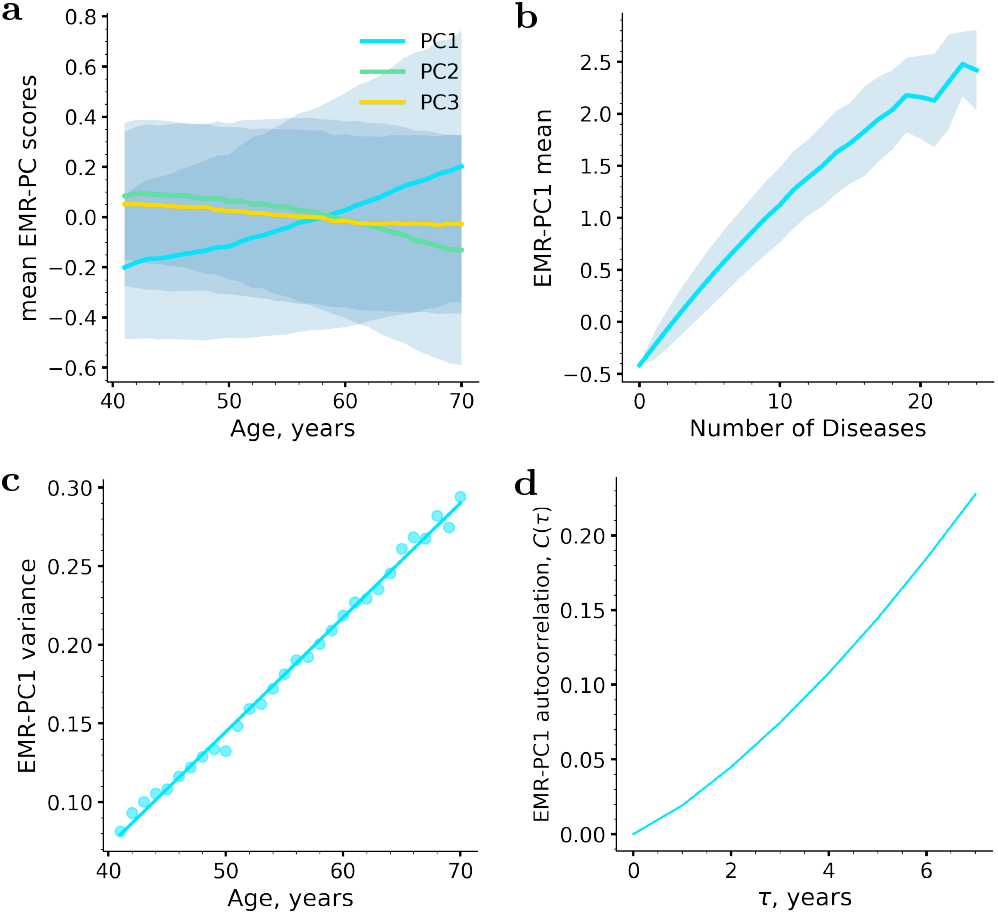
PCA of disease-state vectors from the electronic medical records (EMR) from the UK Biobank: (a) Age dependence of the first three PC scores; (b) PC1 correlates with the total number of chronic diseases; The solid lines and the ranges represent the means and the standard deviations in subsequent age-matched cohorts. (c) Variance of PC1 increases linearly with age (d) Autocorrelation function *C*(*τ*) of PC1 increases linearly as a function of the time lag *τ*.

The dominant aging signature, the first PC in the UKB EMR data (EMR-PC1), evolves approximately linearly as a function of age and is linearly associated with the total number of diagnosed diseases (Fig. 2b). Hence, in line with the results of our DNAm analysis above, the first PC correlates with the total number of depolarization transitions (this time being equal to the disease burden at the time of measurement).

As expected, the variance of EMR-PC1 increases linearly with age (Fig. 2c), which is a hallmark of a stochastic process. This time, however, due to the longitudinal nature of the EMR dataset, we can make a stronger claim by computing the autocorrelation function of EMR-PC1. We observe that the autocorrelator increases linearly as a function of the time lag between the observations, which is typical for a result of a stochastic process with a drift (Fig. 2d, see discussion in the section below).

### B. Aging in a complex regulatory network

To explain the key features of aging signatures, let us consider an organism as a network of interacting functional units (FU). Each of the units can be observed in multiple states of varying physiological capacity. We have already presented examples of such microscopic states corresponding to the different methylation levels of particular CpGs or disease states. However, the language may be used to describe other situations involving, e.g., mutations or conformation changes in biomolecules.

For any given FU *i*, we will focus on the two most-occupied microscopic states (Fig. 3a) corresponding to two adjacent potential wells in the free energy landscape shaped by regulatory interactions. We encode the pair of states by a binary variable *σ*_*i*_ taking values of *σ*_*i*_ = 0 and *σ*_*i*_ = 1, respectively. At the end of development, most FUs are polarized, so that most of the subjects occupy one of the selected states.

**FIG. 3.**
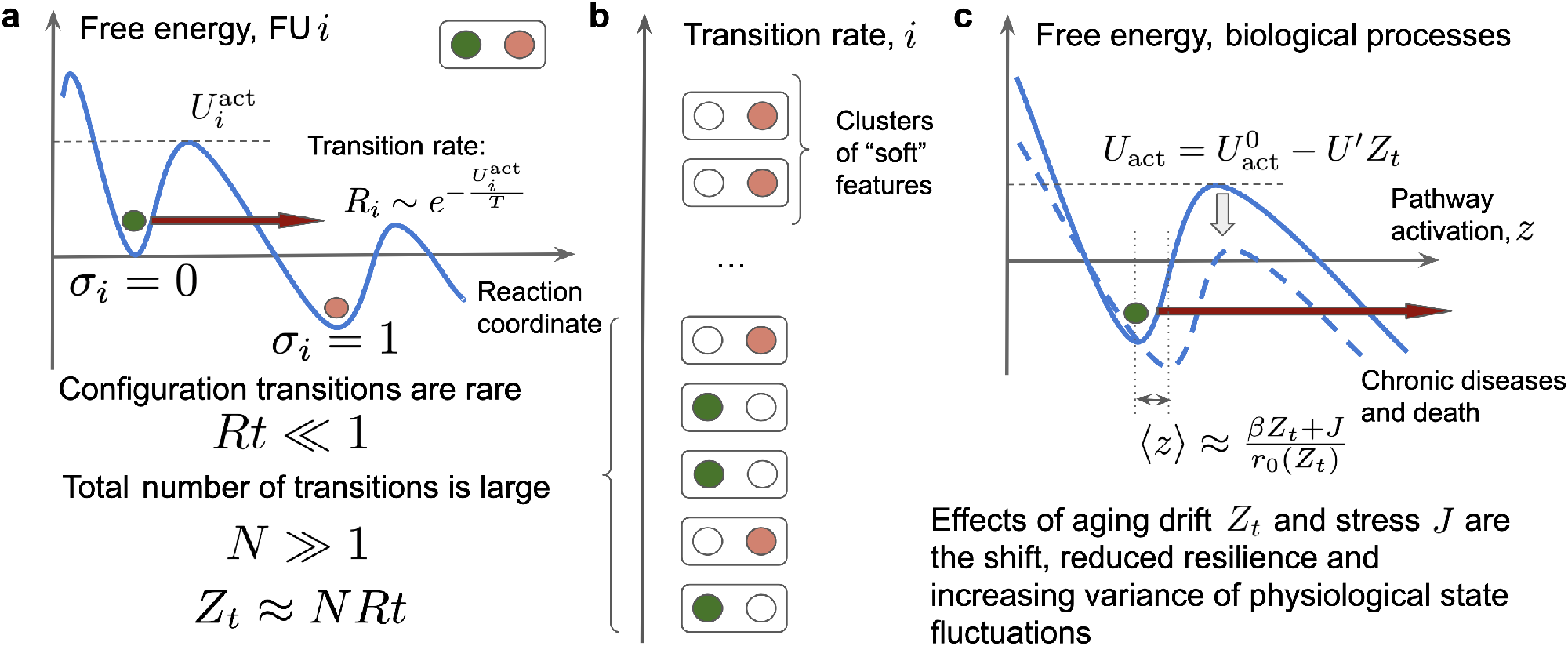
Basic features of the proposed aging model: (a) Schematic representation of relaxation dynamics of a functional unit (FU) *i* residing in a potential well (the blue curve). The subsequent metastable states are labeled by the “polarization” *σ*_*i*_ = 0 and *σ*_*i*_ = 1 (the red arrows indicates thermally activated configuration transition between the microscopic states). The initial, “polarized” state is protected by the activation barrier characterized by the activation energy 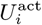. The configuration transition rates *R*_*i*_ are presumed small and depend exponentially on the effective temperature *T*. (b) Human organism consists of a macroscopically large number *N* of FUs. We classify them according to the mean activation rates *R*_*i*_. Most configuration transitions are very rare 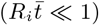. (c) Dynamics of the “soft” FUs with low activation barriers 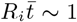. Aging drift causes the reduced resilience and diverging variance of physiological state fluctuations, all proportional to the overall number of the configuration transitions to date *Z*_*t*_ ≈ *NRt*.

Initially polarized states are not necessarily ground states. Hence, over time, the organism state relaxes towards thermal equilibrium via a series of configuration transitions – fluctuations drive the depolarization transitions between the microscopic states. Both the DNAm and EMR data suggest that, in most cases, the configuration transitions are rare: on average, we observe fewer than a single transition between the states over the lifetime 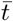 of an organism. In other words, the corresponding transition rates *R*_*i*_ are very small (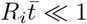, see Fig. 3b).

The low transition rates indicate that the relevant activation energies 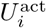 in the free energy landscape defining the states of FUs exceed the effective temperature *T* greatly and are not affected by the effects of aging. Accordingly, we expect that the depolarization rates, 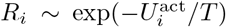, are exponentially small and age-independent.

The effective temperature *T* characterizes the statistical properties of regulatory noise, which may depend on the fidelity of regulatory interactions and the deleteriousness of environment [31]. The effective temperature shall not be confused with (although maybe related to) the body or environmental temperature (see Section S.V).

Quantitatively, stochastic forces cause configuration transitions so that the average polarization of every FU changes linearly over time *t*:

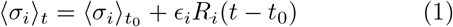

Here the averaging ⟨…⟩_*t*_ occurs over the samples produced at age *t* and *ϵ*_*i*_ = ±1 is the direction of a depolarization transition.

The PCA of a dataset modeled by Eq. 1, would produce the first PC, which is directly proportional to the total number of configuration transitions *Z*_*t*_ ≈*NRt*, where *R* is the characteristic (average) depolarization rate, and *N* is the total number of FUs. It is essential to understand that the total number of FUs jointly describing the organism state, *N*, is practically infinite (*N* ≫1) and hence only a small subset of all FUs may, in principle, be observed directly in any given experiment. We may expect, however, that the total number of the depolarization transitions in any sufficiently large subset of the data (such as DNAm or EMRs) is proportional to *Z*_*t*_.

The depolarization probability for each FU may be small. However, the total number of FUs available for the configuration transitions is huge, and their compound effect does not need to be small. Suppose the lifetime of an organism is sufficiently large. In that case, the total number of configuration transitions *Z*_*t*_ is also very large, and the aging signature described by Eq. 1 should dominate the variance in real-life biomedical data. This is exactly what happens in the PCA of both the DNA methylation (Fig. 1a) and EMR data (Fig. 2a) above. The distribution of the first PC vector’s components would be bi-modal according to the corresponding values of depolarization transition directions *ϵ*_*i*_ (Fig. 1e).

Under the same conditions, *Z*_*t*_ acquire the properties of a random quantity, obeying a stochastic Langevin (or diffusion) equation with a drift. This means that the variance of *Z*_*t*_ (and hence the first PC in the data) in age-adjusted bins should increase linearly with age. This was indeed observed in the DNA methylation (Fig. 1b) and EMR (Fig. 2c) data, respectively.

More evidence in favor of the stochastic character of *Z*_*t*_ (and hence of the first PC scores in the data) could be produced by the investigation of the autocorrelation function *C*(*τ*) = ⟨(*Z*_*t*+*τ*_ *Z*_*t*_)^2^⟩, where ⟨…⟩ stands for the averaging, first, along the individual trajectory and, then over all patients. The autocorrelation function of the leading PC in the EMR data increased linearly as a function of the time lag in the range between 2 and 10 years (Fig. 2d). The diffusion coefficient’s estimates from the variance and autocorrelation increase turned out to be close: 0.012 and 0.009 per year, respectively, thus confirming the association of the leading PC score with the increasing number of configuration transitions *Z*_*t*_.

To highlight the stochastic character of the aging drift in the model, let us note the relation between the number of depolarization transitions *Z*_*t*_ and the configuration entropy in the aging organism. Assuming that we start from highly polarized states, ⟨*σ*_*i*_⟩_*t*_ ≈ 1 −*R*_*i*_*t*, we find that

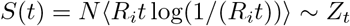

up to a proportionality coefficient. As expected, the configuration entropy increases along with the number of depolarization transitions understood as DNAm changes or the incidence of chronic diseases or age (Figs. 4a and b, respectively).

**FIG. 4.**
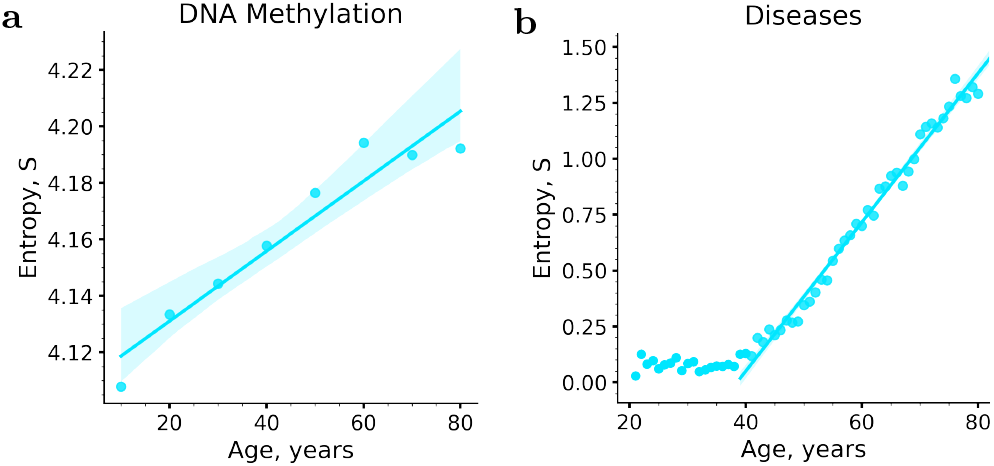
Configuration entropy computed from the distributions of (a) the DNA methylation and (b) EMRs features in subsequent age-matched cohorts increases linearly as a function of age after 40 years.

Let us now turn to the transitions between the microscopic states separated by the smallest activation barriers and hence characterized by the highest depolarization rates (the top FUs in Fig. 3b). In Section S.V, we explain that the interactions between such FUs can no longer be neglected and therefore the FUs should form clusters of co-regulated features. Formally, we expect that the joint activation of FUs forming a cluster (or a pathway) labeled by *A* affects all other FUs *i* in the cluster via a shift of regulatory fields according to 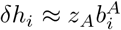 where *z*_*A*_ and 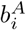 are the pathway activation strength and the participation vectors’ components, respectively.

In Fig. 3c, the solid blue line represents the crosssectional view of the free energy as a function of the pathway activation variable *z*_*A*_ experiencing stochastic fluctuations in response to stress factors. The dynamics of the pathway activation depend on the power of stochastic noise (which is, in turn, proportional to the effective temperature *T*) and persistent stress factors *J*_*A*_. The effects of the regulatory interactions can be described by the recovery rate, *r*_*A*_, which is directly related to the curvature of the basin of attraction for *z*_*A*_. The recovery rate is the inverse recovery time and characterizes the pathway’s ability to respond to stress and relax toward the equilibrium position after a shock.

The configuration transitions occur independently from pathway activations. Each depolarization event exerts regulatory influence and reshapes the free energy landscape for all the other FUs (see the blue dashed line in Fig. 3c). Since the total number of the transitions is very large, the central limit theorem [32] ensures that the net effect of configuration changes on any physiological process must be proportional to the total number of depolarization transitions *Z*_*t*_.

Over longer time scales, well exceeding pathway equilibration times ~ 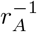 the stochastic component of *z*_*A*_ fluctuations averages out. The mean pathway activation and variance are given by:

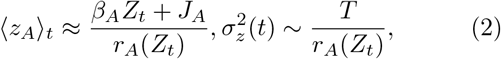

respectively (see Section S.V for the details of the derivation). Here 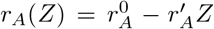 is the age-adjusted recovery rate, whereas *β*_*A*_, and 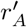 are small and pathwayspecific quantities characterizing the weak mode-coupling effects leading to the compound (and hence proportional to *Z*_*t*_) effect of depolarization processes on pathway activation and resilience, respectively.

Accordingly, the fluctuations of the organism state variables other than those described by Eq. 1 can be attributed to a few clusters of co-regulated features participating in pathways characterized by the smallest recovery rates (vanishing denominators in Eq. 2). In this case, the participation vectors 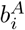 and the pathway activation variables *z*_*A*_ should approximately coincide with the leading PC loading vectors and scores, respectively.

According to Eqs. 2, an increasing number of depolarization transitions *Z*_*t*_ causes progressive shifts in pathway activation. Notably, this effect is indistinguishable, albeit smaller than the effects of constant stress modeled by *J*_*A*_. More subtly, aging in the form of progressive depolarization of an organism state measured by *Z*_*t*_ also affects the recovery rates in the denominator of Eq. 2. The two effects combine and cause the mean pathway activations and therefore the leading PC scores in the data depend on age in a non-linear — hyperbolic fashion (see the dynamics of DNAm-PC3 in Fig. 1c and EMR-PC2 in Fig. 2a).

The non-linear coupling of organism-state fluctuations with depolarization transitions may reduce one of the smallest recovery rates to zero: *r*(*Z*_*t*_) = *r*^0^ *r*′*Z*_*t*_ = *r*^0^(1 − *t/t*_max_) at some point late in life at age *t*_max_ = *r*^0^*/*(*r*′*dZ*_*t*_*/dt*). The situation corresponds to the critical point corresponding to the complete loss of resilience, that is, the inability of the system to retain its homeostasis equilibrium and hence it is incompatible with survival [17].

There is no way to measure the recovery rate in crosssectional data. However, according to Eq. 2, the vanishing recovery rate should lead to the simultaneous divergence of one of the leading PC scores and its variance at a certain advanced age. In our analysis, DNAm-PC3 increases faster than linearly as a function of chronological age. The fit of the DNAm-PC3 scores to the hyperbolic solution for the average *z*_*A*_ from Eq. 2 gives *t*_max_ ≈130 years (see the solid line in Fig. 1c and Section S.I for the details of the calculations).

In agreement with Eq. 2, the extrapolation shows that the inverse variance of DNAm-PC3 hits zero and hence the variance of DNAm-PC3 diverges at approximately 120 years (see the solid line in Fig. 1d). The estimations of the limiting age from the behavior of DNAm-PC3 mean and its variance are comfortably close. Hence, our calculations support the existence of a critical point in the age range of 100 −150 years.

In reality, the disintegration of the organism state happens well before reaching the criticality at the limiting age *t*_max_. Stress factors and the depolarization of the organism state do not merely shift the mean pathway activation levels. Both factors may also decrease the activation energy separating the organism state from disintegration and death (Fig. 3c). In the linear regime, the activation energy linearly depends on the mean-field, 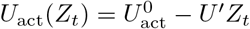 where *U*′ = *dU/dZ*.

The mortality in the model is nothing else but the probability of barrier crossing per unit time: 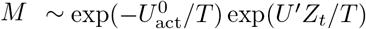. Therefore, the aging drift in the form of configuration transitions registered by the progressively increasing *Z*_*t*_ ~ *t* may drive the exponential acceleration of all-cause mortality with age: *M* ~ exp(Γ*t*). The mortality doubling rate Γ = *T* ^−1^*U* ′*dZ*_*t*_*/dt* in the model depends on the details of the regulatory interactions (through *U′*), the rate of the aging drift *dZ*_*t*_*/dt*, and the effective temperature *T*.

## III. DISCUSSION

We put forward a semi-quantitative model of aging in a complex regulatory network and applied it to the analysis of human aging signatures in a cross-sectional white-blood-cells DNAm dataset [28] and the extensive collection of longitudinal electronic medical records (EMRs) from the UK Biobank [29]. We demonstrated that the key features of the physiological indices dynamics may be explained by the compound effect of massive and thermodynamically irreversible configuration changes accompanied by increasing entropy.

The data suggest that the rates characterizing transitions among the microscopic states of the methylation status or the incidence of specific diseases are small. In most cases, on average, fewer than a single state change occurs throughout lifetime. However, even though the rates of configuration changes may be low, the total number of configuration transitions between the states is vast: no less than 25% of all methylation sites exhibited agerelated dynamics. Hence, the overall number of concurrently occurring transitions is large, so their compound effects dominate the dynamics of the physiological state. We observed that the leading aging signature, which is the first PC score explaining most variance in the data, increased on average linearly with age in the PCA of the DNAm and EMR data. In both cases, the first PC score was proportional to the total number of configuration transitions (the number of DNAm level changes or the total number of chronic diseases). Simultaneously, the variance of the dominant PC score grew linearly with age in both datasets, as is expected for a stochastic quantity, a product of a large number of independent relaxation transitions.

We found that in our examples the total number of configuration transitions *Z*_*t*_, on average, increases linearly with age and explains most of the variance of physiological state variables. Accordingly, we propose using *Z*_*t*_ as a quantitative measure of the net effect of slow configuration changes on the aging organism – thermodynamic biological age (tBA ~ *Z*_*t*_). The dominant aging signature in the data (the first PC score) is then an estimate of tBA from the specific data. Most comfortably, DNAm-PC1 and hence tBA exhibited the strongest correlation to Horvath’s DNAm age.

The total number of transitions is large, and hence tBA increases linearly with age to a very high degree of accuracy (according to the central limit theorem [32]). That may be the mathematical reason why it is almost always possible to build a very accurate predictor of chronological age from different sources of biomedical data (see examples [4, 5, 7]).

Configuration transitions do not only provide a natural clock in aging humans, but also the thermodynamic arrow of time. Our model suggests that tBA is directly related to configuration entropy produced and hence information regarding the healthy state lost in the course of aging. The relationship between tBA and entropy can be understood since the biological age is a single number capturing the result of a large number of independent individual transitions. Particular depolarization patterns may differ in various cells of one subject or among different subjects of the same age. In contrast, the total number of transitions should be similar and hence characterize the overall state of an organism in relation to aging.

Due to limited data availability, we could only analyze aging signatures in DNAm states and chronic diseases. However, we expect that present findings should be universal and generalize to other examples of functional units experiencing configuration changes over time. We may think about (but not limit ourselves to) conformation or chemical modifications of macromolecules, including DNA damage, etc. Since all the configuration changes happen simultaneously and increment tBA, we must not consider any single kind of them as causing the aging drift.

Configuration transitions change the organism’s state and affect all biological processes. Since the number of configuration changes is large, the details of individual transitions are not important. The compound effect of the aging drift manifests itself as a “mean-field” causing the shift of physiological indices that is proportional to the number of configuration transitions to date (and hence to tBA itself).

The mean-field theory is a powerful approximation for understanding the behavior of interacting systems first developed in physical sciences (see, e.g., [33]) and since then applied in statistical inference [34] in general and in specific applications (see, e.g. protein structure prediction [35, 36]). In the present work, we substitute the overwhelming complexity of actual interactions between FUs for a much simpler picture, where each of the FUs or large clusters of FUs operate independently and experience the average effects of the behaviour of all other FUs quantified by tBA.

The mean-field produced by the configuration transitions yields the strongest effects on the large clusters of FUs characterized by long recovery times and hence exhibiting strong fluctuations and dominating the leading PCs other than PC1 ~ tBA in biological signals. We show that the effect of the aging drift on such modules or pathways is similar to the effects of stresses (such as smoking or diet). Since the BA increases linearly with age, we expect that, in the first approximation, all pathways “follow” the aging process by increasing (or decreasing) activations linearly with age.

The proposed model provides a good semi-quantitative explanation how the increasing mean field and the nonlinearity of regulatory interactions produce significant deviations of mean pathway activations from a simple linear age-dependence of non-dominant PC scores in biomedical data (see DNAm-PC3 and EMR-PC2 in the PCA of DNA methylation and EMR data, respectively).

Nonlinear regulatory interactions let the configuration changes (but not stresses) affect the resilience understood as the ability of a cluster of interacting FUs to respond to a perturbation and relax to the equilibrium afterwards. If the recovery rate is particularly small, this may lead to the divergence of organism state fluctuations at some advanced age corresponding to the critical point, where the recovery rate vanishes. This happens with the cluster of DNAm levels associated with the DNAm-PC3. By extrapolation, we observed both the mean and the variance of DNAm-PC3 diverging at the age close to *t*_max_ ≈ 130. Gene set enrichment analysis (GSEA) of genes regu-lated by CpG sites involved in DNAm-PC3 reveals association with innate immunity. Recently, we demonstrated that linear log-mortality predictors built from complete blood counts (CBC) and physical activity [17] also exhibited diverging fluctuations and a vanishing recovery rate at about the same limiting age *t*_max_ ≈130 years. We, therefore, infer that the white-blood-cells DNA methylation, blood composition and even physical activity all substantially depend on a common factor related to innate immunity and all-cause mortality.

The prediction of mortality (or the remaining lifespan) in humans hence requires an estimate for tBA ~ *Z*_*t*_ and for a few most crucial pathway activations (also, on average, depending on *Z*_*t*_). Hence, no single biological age measure fully describes longevity in humans. We expect that the biological age models trained to predict chronological age should yield better estimates of tBA. On the other hand, the models trained to predict the remaining lifespan (such as PhenoAge [8], GrimAge [9], DOSI [17], etc.) should return a combination of pathway activations associated with the prevalence of diseases and accelerated mortality [12] and hence better suited for the detection of reversible effects of diseases, lifestyles and medical interventions [37].

PCA of human data is peculiar since it produces more than a single age-dependent feature. This is not the case in simpler animals such as worms [38], flies [39] or mice [40], where aging could be explained by a simple dynamic instability leading to the exponential disintegration of an organism state [39, 41]. We expect that the entropic contribution to aging has no time to develop in such cases. The biological age is then a dynamic factor, and the effects of aging may be reversible [40].

This work along with the direct dynamics stability analysis of organism state fluctuations in longitudinal biomedical data [15, 17] support the idea that humans (and probably other long-lived mammals, such as naked mole-rats) evolved so that fully grown subjects are metastable until very late in life. The loss of stability is the result of a loss of resilience due to a combined effect of a vast number of independent configuration transitions.

We put forward arguments suggesting that human aging may have a very significant entropic component. Our approach is, therefore, in line with Hayflick’s proposal [42] that distinguishes the genetic determinism of longevity from the stochasticity of the aging process. If the proposition is accurate, we must expect that although the hallmarks of aging (features or activations of specific pathways leading to mortality and morbidity acceleration [1]) can, in principle, be reverted, the expected effects on lifespan may be transient and limited. Any attempts to reduce the dominant aging signature, tBA, would run against the tendency of complex systems to increase their entropy. Any working strategy would require the availability and timely application of an immense number of precise interventions. This is, to say the least, technologically challenging. Accordingly, we must think that aging in humans can be reversed only partially. For example, recent epigenetic reprogramming experiments [43–45] lead to the reversal of epigenetic clock readouts. The fact that the process of aging is entropic does not necessarily mean that one cannot reset some of the organismal subsystems closer to a younger state. The entropic character of aging implies that age-reversal would be limited to a specific organismal subsystem without a full rejuvenation of the whole organism.

Achieving strong and lasting rejuvenation effects in human may thus remain a remote perspective. Our model, however, suggests that there must be a practical way to intercept aging, that is to reduce the rate of aging dramatically. The rates controlling configuration transitions between any two states depend exponentially on the effective temperature. Hence, even minor alterations of the parameter may cause a dramatic drop in the rate of aging. In condensed matter physics, this situation is known as glass transition, where the viscosity and relaxation times may grow by ten to fifteen orders of magnitude in a relatively small temperature range [46–53]. We note, of course, that living organisms are non-equilibrium open systems, and hence the effective temperature must not coincide with body or environment temperatures. Rather, the effective temperature is a measure of deleteriousness of the environment [31].

We speculate that the evolution of long-lived mammals may have provided an example of tuning the effective temperature. Naked mole rats are known for their exceptional stress resistance, DNA repair efficacy [54–57], and translational fidelity [58, 59]. Those factors should reduce noise in regulatory circuits and lower the effective temperature of the system. One example of such tuning may be used to explain the recent studies indicating that naked mole-rats breeders age slower than their non-breeding peers, at least according to the DNAm clock [24].

Social status and mental health also impact the aging rate measured by DNAm and other clocks in humans [60, 61], possibly via neuroendocrine system. Higher socioeconomic status, somewhat counter-intuitively, significantly increases the mortality doubling rate and simultaneously reduces age-independent mortality in such a way that mortality in the highest and lowest income groups converge at an age close to our *t*_max_ estimates [62]. Such a behavior of mortality is consistent with a reduction of the effective temperature in the higher-income cohorts in our model.

Future studies should help establish the best ways to “cool down” the organism state and reduce the rate of aging in humans. The simple linear PCA exemplified here may only help gain a qualitative understanding of underlying processes. We expect that increasing availability of high-quality longitudinal biomedical data will lead to a better understanding of the most critical factors behind the kinetics of aging and diseases, including those controlling entropy production in the course of aging. This should lead to a discovery of actionable targets influencing the rate of aging, help slow down aging and thus produce a dramatic extension of human healthspan.

## IV. ACKNOWLEDGEMENTS

We would like to thank A. Velikanova, D. Kriukov, K. Avchaciov and T. Pyrkov for insightful discussions and help with data preparation; M. Kholin and A. Kadet for stimulating discussions and comments on the manuscript. The work was funded by Gero LLC (Singapore).

## MATERIALS AND METHODS

### S.I. PCA OF THE DNA METHYLATION DATA

We took the white-blood-cells methylation data from GSE87571 dataset [28]. It contains 729 samples (more than 440k features each) collected from patients of both genders (341 males and 388 females) covering the age range between 14 and 94 years of age.

To focus the analysis on aging, we filtered out the patients younger than 20 y.o. (620 samples remaining). We filtered out the CpG sites according to Pearson’s correlation between the DNA methylation levels and the chronological age at the level of *p <* 0.005*/N* (where *N* is the total number of the reported features), thus obtaining 96536 sites. We performed and reported the results of the PCA on the resulting data.

We computed Horvarth’s methylation age as described in [4]. A few CpG sites (cg17099569, cg00431549, cg11025793, and cg14409958) were not present in the data, and hence we had to exclude them from the calculation.

DNAm-PC3 increased with age at a faster than linear pace. We collected all the pairs of the DNAm-PC3 scores and the chronological age for every patient *n* in the dataset and used the available age-range to produce a fit of the data to average from Eq. 2:

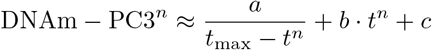

with the uniform Gaussian error and *t*_max_, *a, b* and *c* being the parameters of the fit. The calculation returned *t*_max_ = 129.9 years. We also performed the linear fit of the inverse variance of DNAm-PC3 and obtained 90% CI [114.5, 122.2] for *t*_max_.

**FIG. S.1.**
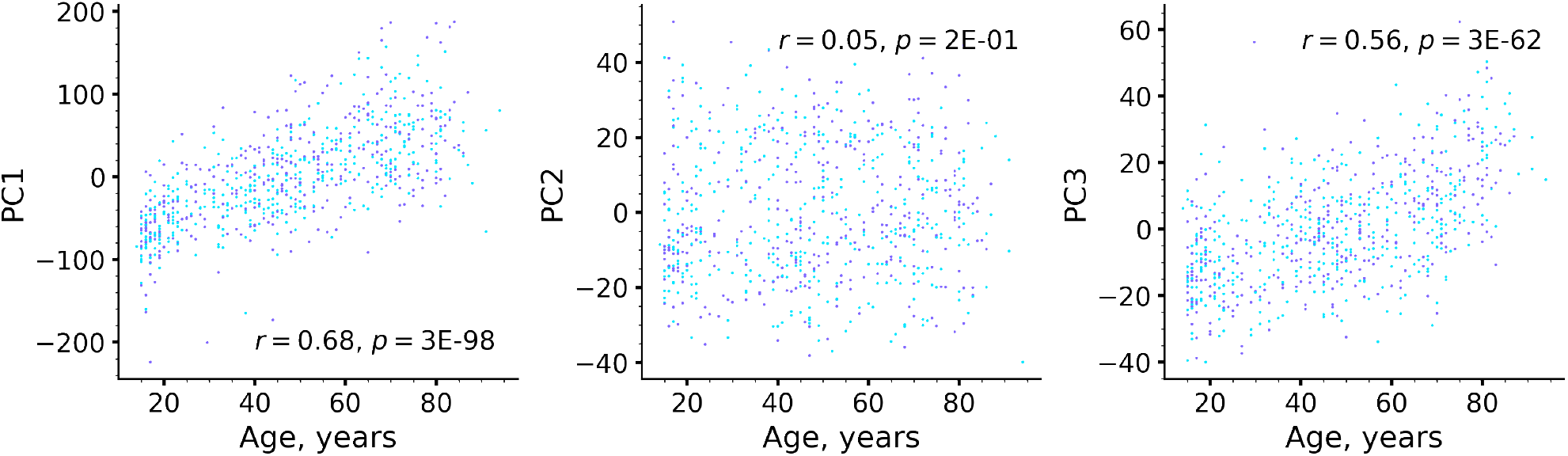
Association of the PC scores in regulatory fields with chronological age.

**FIG. S.2.**
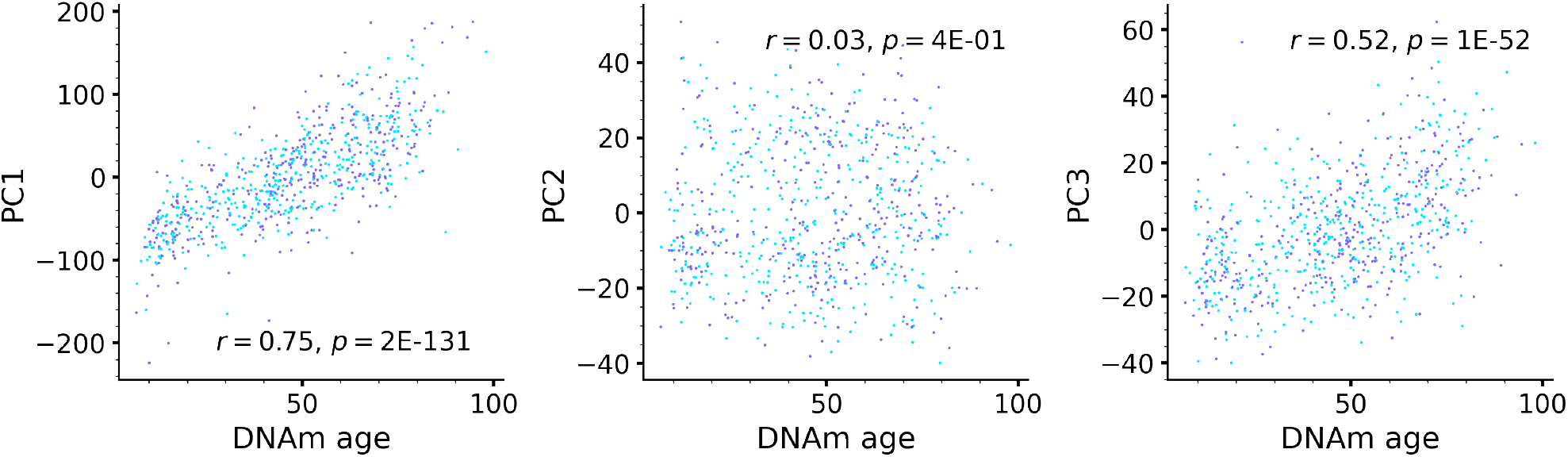
Association of the PC scores in regulatory fields with Horvath’s methylation clock (DNAm age).

### S.II. GENE SET ENRICHMENT ANALYSIS

We collected the CpG sites best associated with DNAm-PC1 and DNAm-PC3 according to the values of the respective vector components. We retrieved the gene IDs from Illumina’s 450k methylation arrays documentation. Finally, we performed Gene Ontology (GO) and disease ontology (DO) enrichment with the help of the R “clusterProfiler4.0” package [63].

### S.III. PRE-PROCESSING OF EMRS FROM UK BIOBANK

To avoid using the disease labels corresponding to the transient diseases, we selected 111 chronic diseases diagnoses using Chronic Condition Indicators for ICD-10 [64]. Overall, in the EMR dataset are 389494 patients, of mostly Caucasian origin (366715 or 94%), of both sexes (179032 males and 210462 females) in the age range 38 − 74).

### S.IV. ENTROPY/ENTROPY PRODUCTION RATE DETERMINATION

For the practical calculation of entropy, we used a Python library *scipy*.*stats*.*entropy* [65], which was applied to the individual distributions of methylation levels and to the distributions of EMR vectors averaged over the population in age-binned cohorts.

### S.V. THEORY: AGING IN A COMPLEX REGULATORY NETWORK

We propose to model the effect of the interactions among FUs with the help of the auxiliary variables – the effective “regulatory fields” *h*_*i*_ evolving over time according to

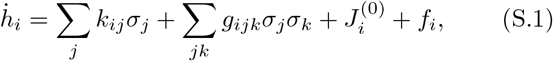

where *k*_*ij*_ and *g*_*ikj*_ describe the first linear and first order non-linear interaction between the individual units. The force terms 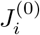 and *f*_*i*_ represent the effects of constant (such as smoking or diets) and stochastic (social status, deleteriousness of the environment [31]) factors, respectively. For simplicity, we assume that the noise factors have zero mean and are not correlated over time. The states of individual FUs *i* can be observed depending on the regulatory field *h*_*i*_ according to the Boltzmann distribution: 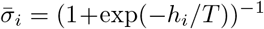 where *T* is the effective temperature.

We start from Eqs. S.1 and observe, that the regulatory fields change over time in response to the deterministic (the direct linear and the higher-order non-linear interactions between the units) and stochastic forces *f*_*i*_. We naturally assume that the stochastic force terms are not correlated over long time intervals: ⟨*f* (*t*)*f* (*t*′)⟩ = *Bδ*(*t* − *t*′) with *B* is the power of the stochastic noise, ⟨…⟩ stands for the averaging along the individual trajectory and over all specimen, and *δ*(*t*) is the Dirac delta-function.

In spite of apparent simplicity, the Eqs. S.1 are non-linear and may have highly non-trivial solutions leading to applications in condensed matter physics [66] and neurophysiology [67]. For our discussion, it is important that the stochastic noise drives the system towards equilibrium at an effective temperature controlled by the power of the noise *T* ~ *B*.

The data suggests that there is a large “bulk” of units characterized by excessive lifetimes. Mechanistically this may be explained by operating within a vicinity of a metastable state with a very high activation energy *U*_act_ relative to the effective temperature, *U*_act_ ≪ *T* (Fig. 3a). We will assume that the effects of aging are small on the scale of *U*_act_ and hence the depolarization rates *R*_*i*_ *U*_act_≫*T* are not only very small, but also do not considerably depend on age. Accordingly, the depolarization is on average a linear function of age and the total number of configuration transitions *Z*_*t*_: ⟨Δ*σ*_*i*_⟩ = *R*_*i*_*t* ∝ *Z*_*t*_ and 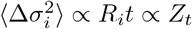.

Let us think that the aging drift in the form of simultaneously occurring configuration transitions progresses slowly compared to fast functional responses in the organism. We linearize the equations for the regulatory fields next to the youthful state 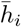:

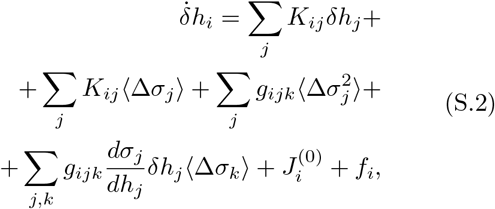

where 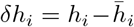 and Δ*σ*_*j*_ variables describe the deviations of the fields and depolarization of the units, whereas the averages ⟨…⟩ involve the averaging over the “bulk” uncorrelated states only.

The solutions of the linearized Eq. S.2 can be best understood with the help of a linear decomposition: 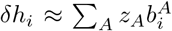 where *z*_*A*_ are the pathway activations, and 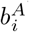 are the right eigenvectors of the interaction matrix corresponding to the smallest eigenvalues *r*_*A*_ (the matrix *K* is non-symmetric and hence its complete eigensystem must include the left, *a*^*A*^*K* = −*r*_*A*_*a*^*A*^, and the right, *Kb*^*A*^ = − *r*_*A*_*b*^*A*^, eigenvectors). The components of the vector 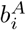 characterize the participation of the FU *i* in the pathway *A*.

Substituting the solution into the equation and multiplying both sides by the corresponding left eigenvector, we find, that

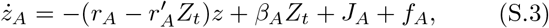

where *J*_*A*_ = *a*^*A*^*J, f*_*A*_ = *a*^*A*^*f*. The effect of aging comes through the mean field on the pathway activation *β*_*A*_*Z*_*t*_ = *a*^*A*^*K* ⟨Δ*σ*⟩ + 𝒪 (*g*) and the non-linear correction to the eigenvalue 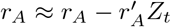.

It is important to understand, that all the relevant vectors and constants can not be derived and could only be measured experimentally. The large number of configuration transitions ensures by virtue of the central limit theorem that the effect of the mean field is exactly linear in *Z*_*t*_.

Qualitatively, the net effect of the rare transitions and the associated mean field *Z*_*t*_ together produce a persistent pathway activation, on average, slowly increasing with age. This is often referred to as an enslavement principle: stochastic depolarization transitions produce a slowly evolving mean field *Z*_*t*_ that disturbs pathways characterized by fast relaxation times having thus enough time to adjust to its current level.

